# Increasing the reliability of functional connectivity by predicting long-scan functional connectivity based on short-scan functional connectivity: model exploration, explanation, validation, and application

**DOI:** 10.1101/2023.06.09.544367

**Authors:** Bo Hu, Ying Yu, Yu-Ting Li, Ke Wu, Xiao-Tian Wang, Lin-Feng Yan, Wen Wang, Guang-Bin Cui

**Affiliations:** Department of Radiology, Functional and Molecular Imaging Key Lab of Shaanxi Province, Tangdu Hospital, Air Force Medical University (Fourth Military Medical University), Xi’an, China; Department of Radiology, Air Force Hospital of Eastern Theater Command, Malu Road, Nanjing, China; School of Information and Communication Engineering, Xi’an Jiaotong University, Xi’an, China

## Abstract

Functional connectivity (FC) is a widely used imaging parameter of functional magnetic resonance imaging (fMRI). However, low reliability has been a concern among researchers, particularly in small-sample-size studies. Previous studies have shown that FC based on longer fMRI scans was more reliable, therefore, a feasible solution is to predict long-scan FCs using existing short-scan FCs. This study explored three different generalized linear models (GLMs) using the human connectome project (HCP) dataset. We found that the GLM based on individual short-scan FC could effectively predict long-scan individual FC value, while GLMs based on whole-brain FCs and dynamic FC performed better in predicting long-scan summed FC value of whole brain. The models were explained through visualization of weights in models. Besides, the differences in three GLMs could be explained as differences in distribution features of FC matrices predicted by them. Results were validated in different datasets, including the Consortium for Reliability and Reproducibility (CoRR) project and our local dataset. These models could be applied to improve the test-retest reliability of FC and to improve the performance of connectome-based predictive models (CPM). In conclusion, we developed three GLMs that could be used to predict long-scan FC from short-scan FC, and these models were robust across different datasets and could be applied to improve the test-retest reliability of FC and the performance of CPM.

## 1. Introduction

Functional connectivity (FC) is a well-known image feature that is measured by the Pearson correlation coefficient of functional magnetic resonance imaging (fMRI) signals between two brain regions (1). FC was effective in diagnosing brain diseases (2), individual identification (3), and exploring neurobiological mechanisms (4). However, researchers found that FC suffered from low reliability, particularly in small-sample-size studies (5, 6). Given that reliability is the foundation of experimental accuracy, there is an urgent need to improve the reliability of FC to enhance the overall quality of FC-related researches.

Previous studies have shown that functional connectivity (FC) based on longer fMRI scans (especially longer than 25 minutes) was more reliable (7, 8). However, due to the significant time and resource consumption, fMRI scans were relatively short (around 6 minutes) in many previous studies (9, 10). To prevent the great waste of existed short-scan fMRI data, several methods was explored to extend its scan length, including concatenating fMRI data from multiple runs (11) or modalities (12). However, these methods are only applicable in cases where multi-run or multimodal data are available. For studies with only a single-run data, an alternative approach is to predict long-scan FCs based on short-scan FCs.

Tavor et al. (13) utilized a linear model to predict the activation pattern of fMRI signals in multiple task states based on the activation pattern in the resting state. This approach extended the application scope of task-state fMRI to individuals who were unable to perform cognitive tasks. Additionally, several studies successfully predicted the structural connectivity (14), brain activity (15), and brain atrophy (16) based on resting state FC. These studies demonstrated the potential of resting-state FC for image-to-image prediction, indicating that short-scan FCs could potentially be used to predict long-scan FCs. However, relevant study has not been conducted yet.

In this study, our objective was to predict long-scan FC from short-scan FC using a four-step process (**Fig 1a**). Firstly, we constructed three different generalized linear models (GLMs) based on the human connectome project (HCP) dataset (17). We found that the GLM based on individual short-scan FC could effectively predict long-scan individual FC value, while GLMs based on whole-brain FCs and dynamic FC performed better in predicting long-scan summed FC value of whole brain. The models were explained through visualization of weights in models. Besides, the differences in three GLMs could be explained as differences in distribution features of FC matrices predicted by them. Thirdly, our results were robust across the Consortium for Reliability and Reproducibility (CoRR) Project (17) and our local dataset (18, 19). Finally, our methods had the potential to enhance the test-retest reliability of FC and the performance of connectome-based predictive models (CPM).

**Fig 1.**
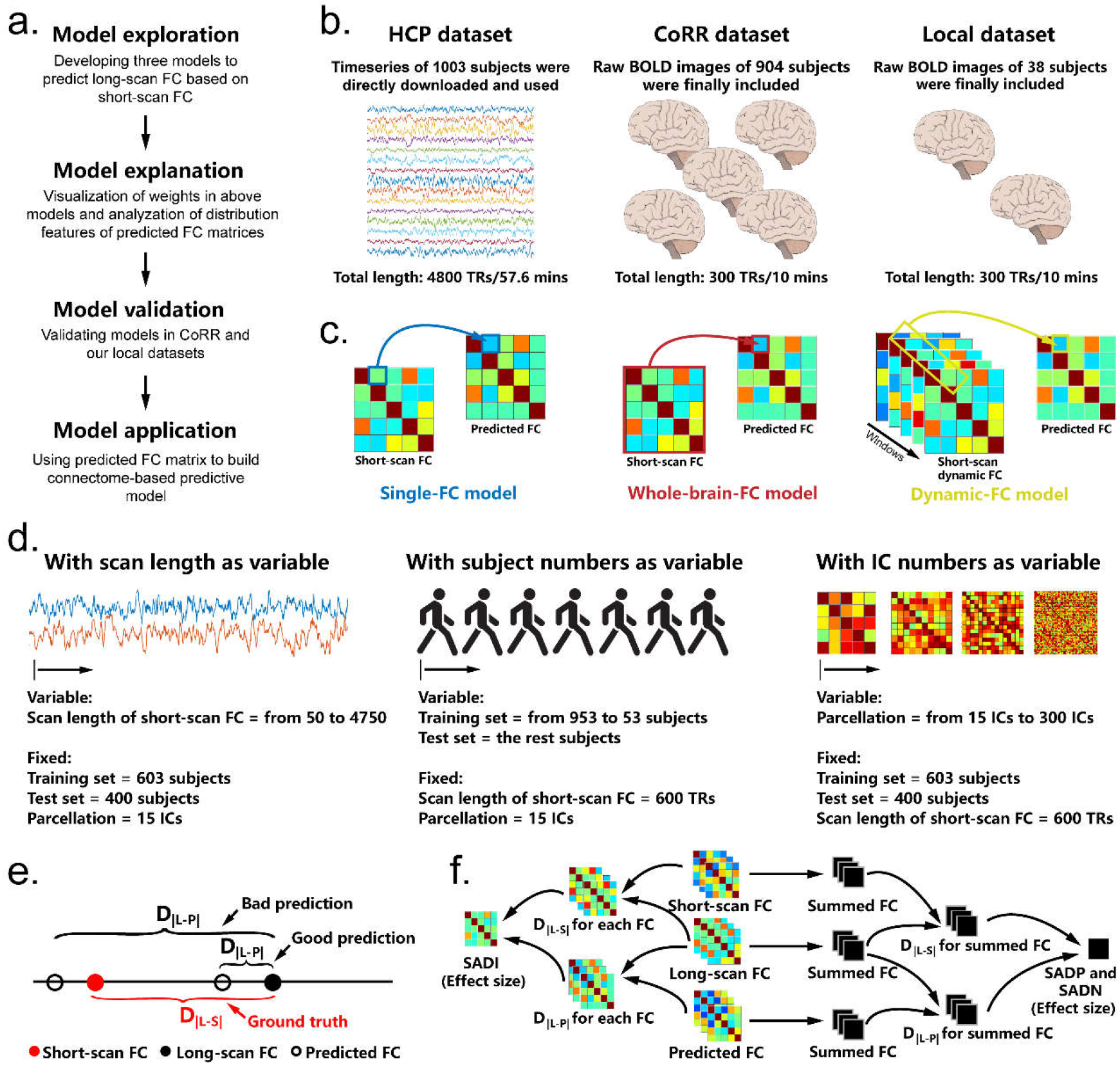
Study design, details of models, and outcome assessments. (a) Four main parts of this study. (b) Datasets used in this research. (c) Three models in this research. For each panel, the matrix in the left was original short-scan FC matrix, and the matrix in the right was the FC matrix that predicted by each model. The first model was single-FC model, which directly used the short-scan FC to predict its long-scan form. The second was whole-brain-FC model, which used all FCs in the brain to predict each long-scan FC. The third was dynamic-FC model, which used a sliding-window method to extract the dynamic FC to predict its long-scan form. (d) Variables in three models. All conditions of each variable were tested, while the other variables were fixed. (e) The definition of absolute distance of FC. The absolute distance D_|L-S|_ was defined as the absolute difference between long-scan FC and short-scan FC, and the absolute distance D_|L-P|_ was defined as the absolute difference between long-scan FC and predicted FC. D_|L-S|_ > D_|L-P|_ mean that the prediction was good, because the predicted FC was more similar to long-scan FC than the short-scan FC, otherwise, it was a bad prediction. (f) Three parameters of absolute FC distance were calculated, including the shortened absolute FC distance of individual FC (SADI), the shortened absolute FC distance of the sum of all positive FCs (SADP), and the shortened absolute FC distance of the sum of all negative FCs (SADN). SADI was measured by calculating the effect size (Cohen’s d) between the D_|L-S|_ and D_|L-P|_ of individual FC. SADP and SADN were obtained by separately summing all positive or negative FCs in the brain and calculating the effect size (Cohen’s d) between the D_|L-S|_ and D_|L-P|_ of the summed FC.

## 2. Materials and methods

### 2.1 Subjects and imaging data

Details of demographic information, scanning parameters and data processing procedures were displayed in **Supplementary methods** and **Table S1**.

#### 2.1.1 HCP dataset

The HCP young adults dataset (20) (**Fig 1b**) consisted of 1003 subjects between the ages of 22 and 35. The timeseries of all brain nodes from all subjects were directly downloaded and used (https://db.humanconnectome.org/data/projects/HCP_1200), which were extracted from the minimally pre-processed fMRI data (21). Each subject completed four runs of 14.4-minute resting-state fMRI scans with a 0.72-second time of repetitions (TR), resulting in a total scan length of 4800 TRs (or frames) (4 runs×14.4 min/run×60 s/min×1/0.72 TR/s). The whole brain was parcellated into different independent components (ICs) using group independent component analysis, resulting in 15, 25, 50, 100, 200, or 300 ICs. This resulted in 1003 (subjects)×4800 (scan length)×N (ICs) timeseries.

#### 2.1.2 CoRR dataset

To ensure high quality of the study, 904 subjects with the same TR (2 seconds) were selected from 18 sites from the CoRR dataset (17). All participants were between the ages of 7 and 84 and completed at least 2 runs of fMRI scan, with each run lasting at least 5 minutes, resulting in a total scan length of at least 300 TRs (2 runs×5 min/run×60 s/min×1/2 TR/s). A traditional pipeline of pre-processing and denoising was used through the Statistical Parameter Mapping and DPABI toolkit (22). The region of interest (ROI) was defined as the coordinates of peak values of ICs in the HCP dataset (15-IC parcellation), and time series were extracted by averaging values around the ROI (radius = 6 mm). Finally, 904×300×15 timeseries were obtained.

#### 2.1.3 Local dataset

Our Local dataset (18, 19) included 38 subjects that aged between 18 and 22, and all subjects completed 2 runs of fMRI scan (5 mins per run, TR = 2 s). Finally, 38×300×15 timeseries were obtained.

### 2.2 Model exploration: developing three models to predict long-scan FC based on short-scan FC

#### 2.2.1 Models

The study employed three different GLMs (**Fig 1c**) to predict the long-scan FC. The first model, known as the single-FC model, used the individual short-scan FC as independent variable. The second model, known as the whole-brain-FC model, used all short-scan FCs in the brain as independent variables. The third model, known as the dynamic-FC model, used a sliding-window method to extract the dynamic FC as independent variables. Dynamic-FC model used a common window width (40 TRs) and step length (3 TRs) in extracting dynamic FC, although other conditions were also tested.

#### 2.2.2 Variables

Three variables may influence the result, including scan length, subject numbers, and methods of parcellation. Consequently, all conditions of each variable were tested while the condition of other two variables were fixed at the same time (**Fig 1d**).

##### With scan length as variable

Long-scan FC were generated by correlating long-scan fMRI timeseries (4800 TRs). Short-scan FC was calculated by extending the scan length from 50 TRs to 4750 TRs, with an interval of 50 TRs. The brain was parcellated into 15 ICs, and 603 subjects (∼60%) were randomly selected as the training set, while the remaining 400 subjects (∼40%) were used as the test set.

##### With subject numbers as variable

The training set were successively reduced from 953 subjects to 53 with an interval of 50 subjects, and the rest subjects were the test set. To avoid bias in subject selection, the selection process was randomly repeated 500 times and the results were presented as the mean ± standard deviation of 500 repetitions. Short scan was defined as fMRI timeseries within the first half of the first run (600 TRs, approximately 7.2 minutes) to align with the common scan time. Additionally, the brain was parcellated into 15 ICs.

##### With independent component numbers as variable

The brain was parcellated into different ICs (from 15 ICs to 300 ICs). Short scan was defined as fMRI timeseries within the first half of the first run, and 603 subjects were randomly selected as the training set and the rest 400 subjects as the test set.

#### 2.2.3 Outcome assessments

To evaluate the results, we compared the absolute distance (**Fig 1e**) between long-scan FC and short-scan FC (D_|L-S|_) with the absolute distance between long-scan FC and predicted FC (D_|L-P|_). The efficacy of the model was determined by assessing if the long-scan FC was closer to the predicted FC than to the short-scan FC (D_|L-P|_< D_|L-S|_).

### 2.3 Model explanation: visualization of weights in above models and analyzation of differences between models

The weights in above three models were visualized to explain the contribution of each component in each model. Besides, the distribution features of FC matrices predicted by above three models were compared, including mean value, standard deviation, kurtosis, and skewness, between the three models.

### 2.4 Model validation: validating models in CoRR and our local datasets

The long-scan fMRI timeseries consisting of 300 TRs were correlated to create the long-scan FC matrix. The short-scan FC was calculated by gradually increasing the scan length from 40 TRs to 290 TRs at intervals of 10 TRs. The brain was divided into 15 ICs. The subjects were randomly divided into a training set (565 subjects, 60%) and a test set (377 subjects, 40%). In the second step, a leave-one-site-out cross validation method was used where each set was selected as the test set and the rest were used as the training set.

### 2.5 Model application: applying predicted FC matrix on the test-retest reliability and connectome-based predictive model

In the first step, the test-retest reliability was measured among the original FC values based on four runs of fMRI scan and among the predicted FC values. The ratio of increase of test-retest reliability was then calculated.

In the second step, the FC matrices (to obtain best performance, the brain was parcellated into 300 ICs) of 400 subjects in test set were used to predict language ability (including oral reading recognition and picture vocabulary), cognitive ability (including fluid, crystal, and total cognitive function composite), and working memory (N-back task).

### 2.6 Statistical analysis

#### 2.6.1 Model exploration

##### The shortened absolute FC distance of individual FC (SADI)

Effect size (Cohen’s d) was calculated to qualify the shortened absolute FC distance from D_|L-S|_ to D_|L-P|_ (**Fig 1f**).

##### The shortened absolute FC distance of the sum of all positive FCs (SADP)

All positive values of the short-scan, long-scan, and predicted FC matrices were respectively summed to generate a value (**Fig 1f**). Then, D_|L-S|_ was calculated by the absolute difference between the summed value of short-scan FC and the summed value of long-scan FC, and D_|L-P|_ was calculated by the absolute difference between the summed value of predicted FC and the summed value of long-scan FCs.

##### The shortened absolute FC distance of the sum of all negative FCs (SADN)

Similar to above, but all negative values in FC matrices were summed together.

#### 2.6.2 Model explanation

Analysis of variance (ANOVA) or the non-parametric Mann-Whitney U test was utilized to compare the features of distribution between short-scan, long-scan, and predicted FC matrices, depending on their adherence to Gaussian distribution.

#### 2.6.3 Model validation

The same as *Model exploration* step.

#### 2.6.4 Model application

##### Test-retest reliability

The intraclass correlation coefficient (ICC) of four runs of FC was calculated to represent the test-retest reliability.

#### Efficacy of CPM

Leave-one-out cross-validation was used for CPM training and test (32). When all procedures were completed, there was a predicted value for each subject, and correlation analysis between predicted values and true values were used to reflect the efficacy of CPMs. The comparison of correlation coefficients were conducted by using Hittner’s *Z* test (23) on cocor toolbox (24), and Bonferroni method was used for multiple comparison correction.

## 3. Results

### 3.1 Model exploration: three models can predict long-scan FC based on short-scan FC with medium effect

#### 3.1.1 With scan length as variable

The shorter the scan length, the higher the SADI for the single-FC model (**Fig 2a**). Additionally, the mean SADI was consistently high for the single-FC model across all scan lengths (**Fig 2b**). However, when the scan length was short, the SADP and SADN of the whole-brain-FC model were better than those of the single-FC model. Furthermore, the dynamic-FC model outperformed the other two models in certain scan lengths in terms of SADP and SADN. The details of results of single-FC models were showed in **Fig 2c and 2d**.

**Fig 2.**
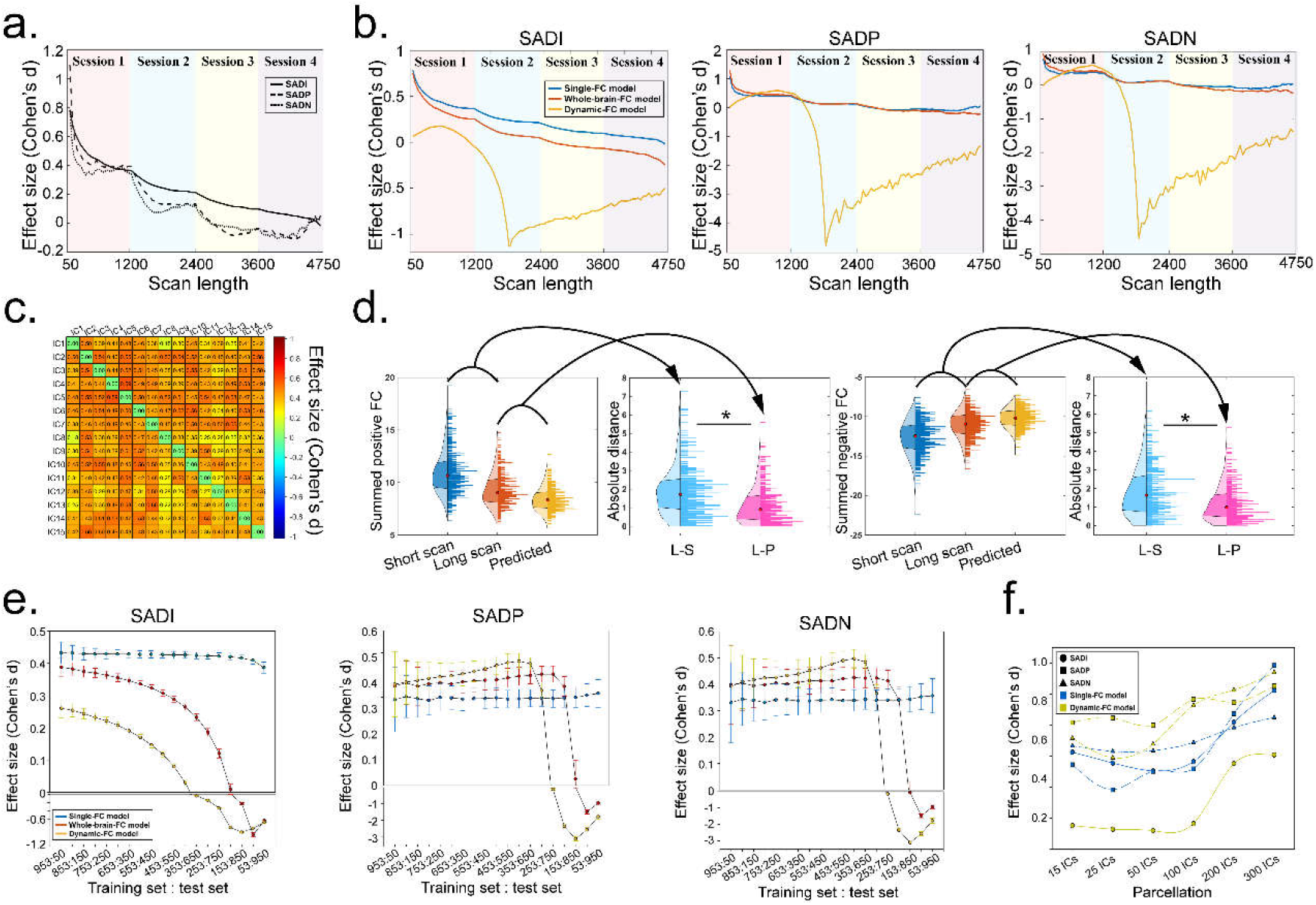
Results of model exploration. (a) The single-FC model showed similar trend across all scan lengths, that is, the shorter the scan length, the more shortened absolute FC distance of a individual FC (SADI). (b) SADI, shortened absolute FC distance of the sum of all positive FCs (SADP) and shortened absolute FC distance of the sum of all negative FCs (SADN) of three models across all scan lengths. (c) Details of SADI of all FCs that predicted by single-FC model (Scan length = 600 TRs; Subjects in training set = 603, subjects in test set = 400; IC numbers = 15). (d) Summed FC values and the comparison of absolute FC distance (variables were the same as above). (e) SADI, SADP, and SADN of three models across all ratios of subjects between training set and test set. (f) SADI, SADP, and SADN of three models across all methods of parcellation.

#### 3.1.2 With data size as variable

The influence of data size on the single-FC model was small (**Fig 2e**). The mean SADI was consistently high in the single-FC model across all data sizes. However, when the training set-test set ratio was higher than 353:650, the SADP and SADN of the whole-brain-FC model and dynamic-FC model outperformed the single-FC model.

#### 3.1.3 With numbers of IC as variable

As the number of ICs increased, both the single-FC model and dynamic-FC model showed improved performance (**Fig 2f**). However, the whole-brain-FC model performed poorly due to an excess of FCs compared to subjects, resulting in a rank deficiency in the GLM model. While the single-FC model had a higher SADI, the dynamic-FC model had higher SADP and SADN.

### 3.2 Model explanation: visualization of weights in above models, and the distribution features of predicted FC matrices were different between models

All long-scan FCs were strongly correlated with short-scan FCs (mean *r* = 0.695). This explains why the single-FC model performed well, as shown in **Fig 3a**. However, when it comes to the whole-brain-FC model, while the highest weight for each long-scan FC was from its short-scan form, other FCs also contributed (as seen in **Fig 3b**, and **S1**). Additionally, the weight of each FC window in the dynamic-FC model fluctuated (**Fig 3c)**, which can be attributed to the inherent properties of fluctuation of FC (25). We found that the mean SADI in the single-FC model was higher than the whole-brain-FC model and dynamic-FC model, however, the SADP and SADN were lower. Consequently, we investigated the reason for the tradeoff between individual FC and summed FC and discovered that the FC matrices predicted by the single-FC model were more centralized, while those predicted by the other two models were more discrete (**Fig 3d-3g**). This was evidenced by decreased standard deviation (P<0.001) and increased kurtosis (P=0.072) in FC matrices predicted by the single-FC model. The tradeoff between the models, outcomes, and FC distributions were further explored in **Fig S2**.

**Fig 3.**
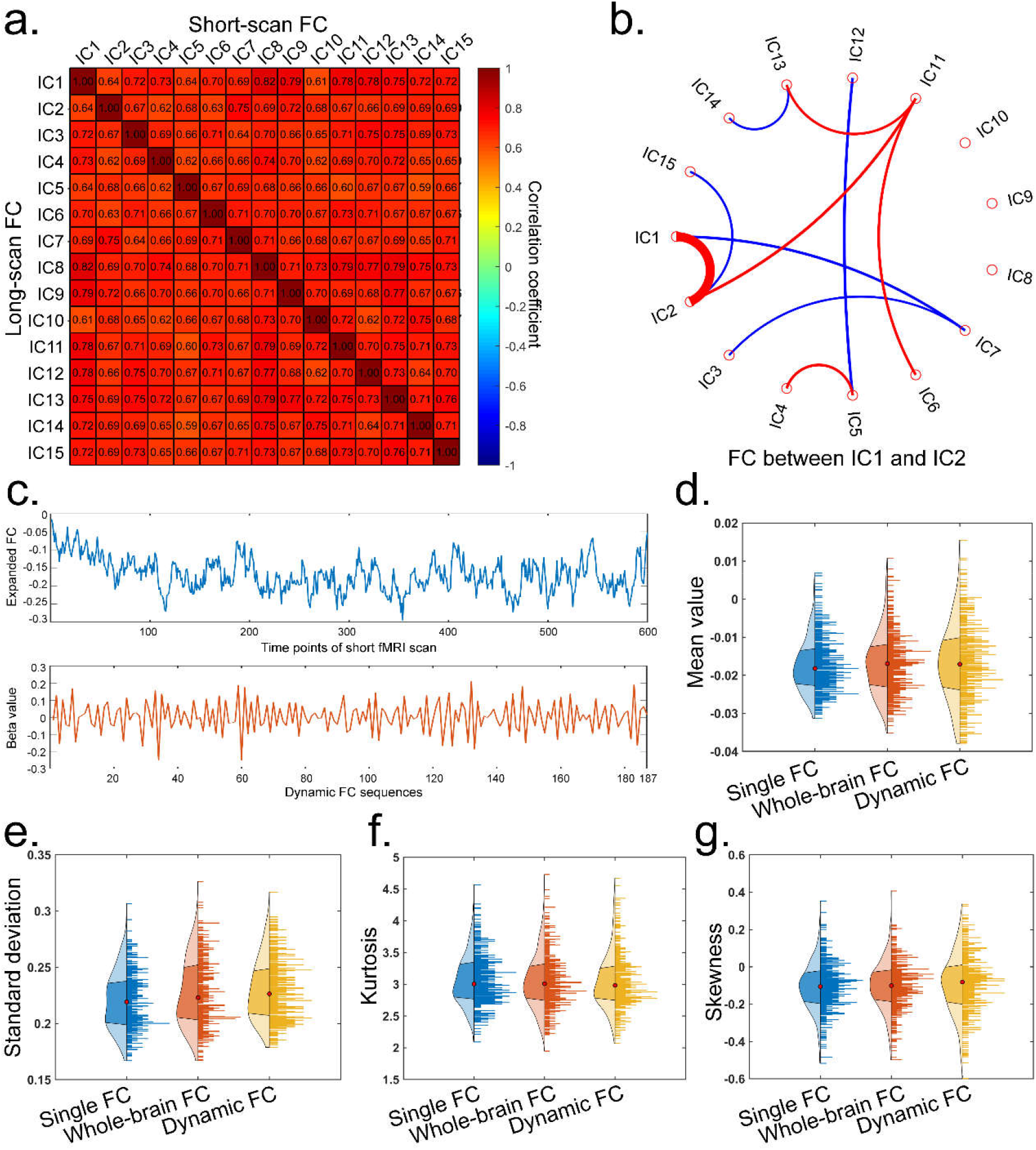
Visualization of weights in above models and analyzation of distribution features of predicted FC matrices. (a) The correlation coefficients between short-scan FCs and long-scan FCs. (b) The strongest weights (top 10) in the whole-brain-FC model to predict the long-scan FC between IC1 and IC2 (others were displayed in **Fig S1**). (c) The weights of dynamic-FC model and the corresponding values of expanded FC across timeseries. (d) The comparison of mean value of FC matrices between three models. (e) The comparison of standard deviation of FC matrices between three models. (f) The comparison of kurtosis of FC matrices between three models. (g) The comparison of skewness of FC matrices between three models.

### 3.3 Model validation: the performances of all models were robust in different data source

The results obtained from CoRR and local datasets were consistent with those from the HCP dataset. Detailly speaking, the single-FC model had a higher SADI compared to the other two models (**Fig 4a**). However, when the scan length was short, the whole-brain-FC model performed better than the single-FC model in terms of SADP and SADN (**Fig 4b and 4c**). In addition, the dynamic-FC model outperformed the other two models in certain scan lengths.

**Fig 4.**
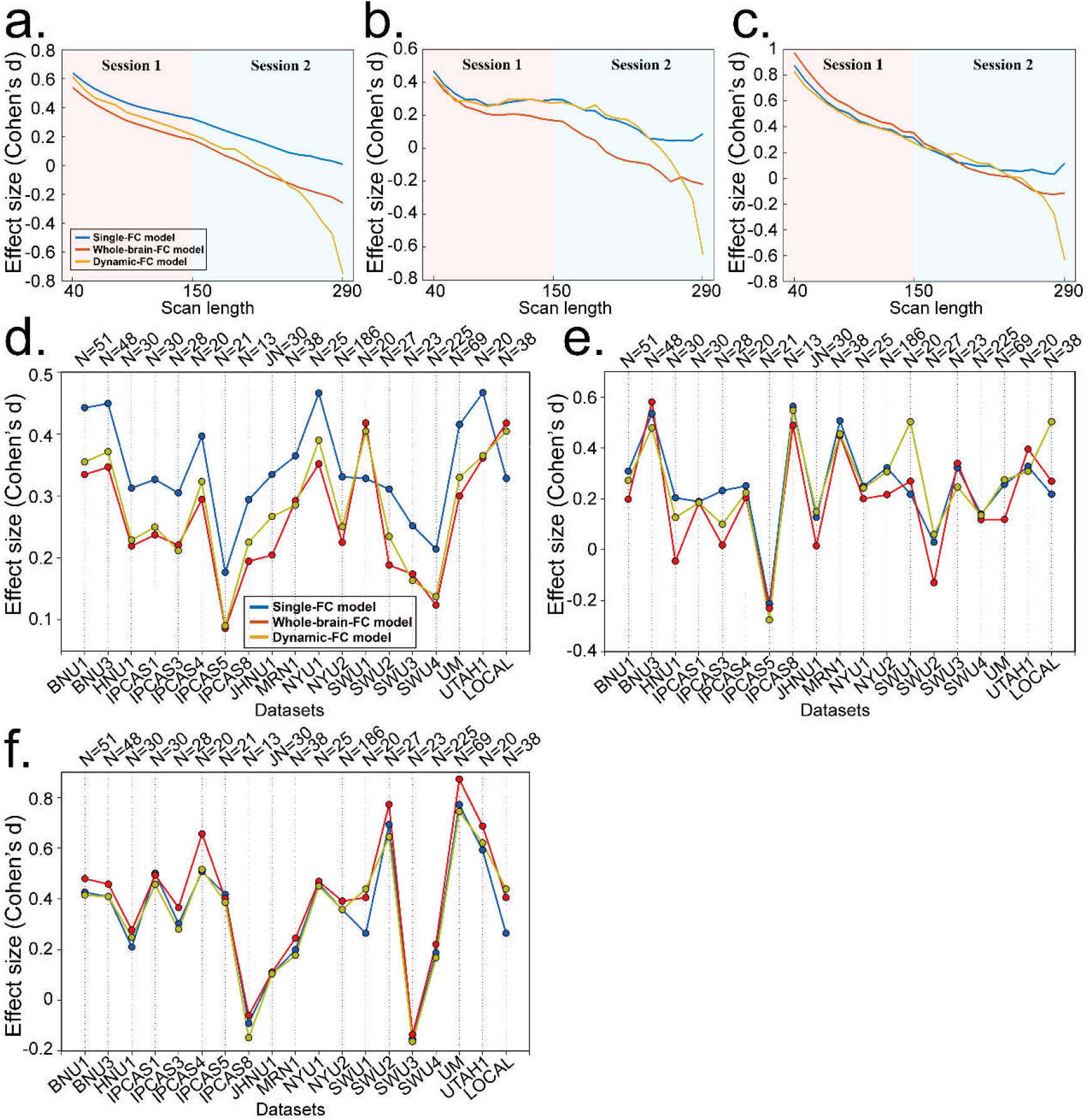
The validation of models on CoRR and our local datasets. (a) The validation of SADI, SADP, and SADN of three models across all scan lengths by using CoRR and our local datasets. (b) A leave-one-site-out method to validate the SADI, SADP, and SADN of three models.

However, the results of the leave-one-site-out cross-validation varied significantly across different sites. The mean SADI ranged from 0.09 (Cohen’s d) to 0.47 (**Fig 4d)**, while the SADP (**Fig 4e)** and SADN (**Fig 4f)** ranged from -0.28 to 0.58 and -0.16 to 0.87, respectively.

### 3.4 Model application: our methods improved the test-retest reliability and the performance of CPM

For both individual FC and summed FC, the ICC of predicted FC was higher than the original FC (**Fig 5a-5c)**. The models performed well when the scan length was shorter than 250 TRs, with the single-FC model performing the best.

**Fig 5.**
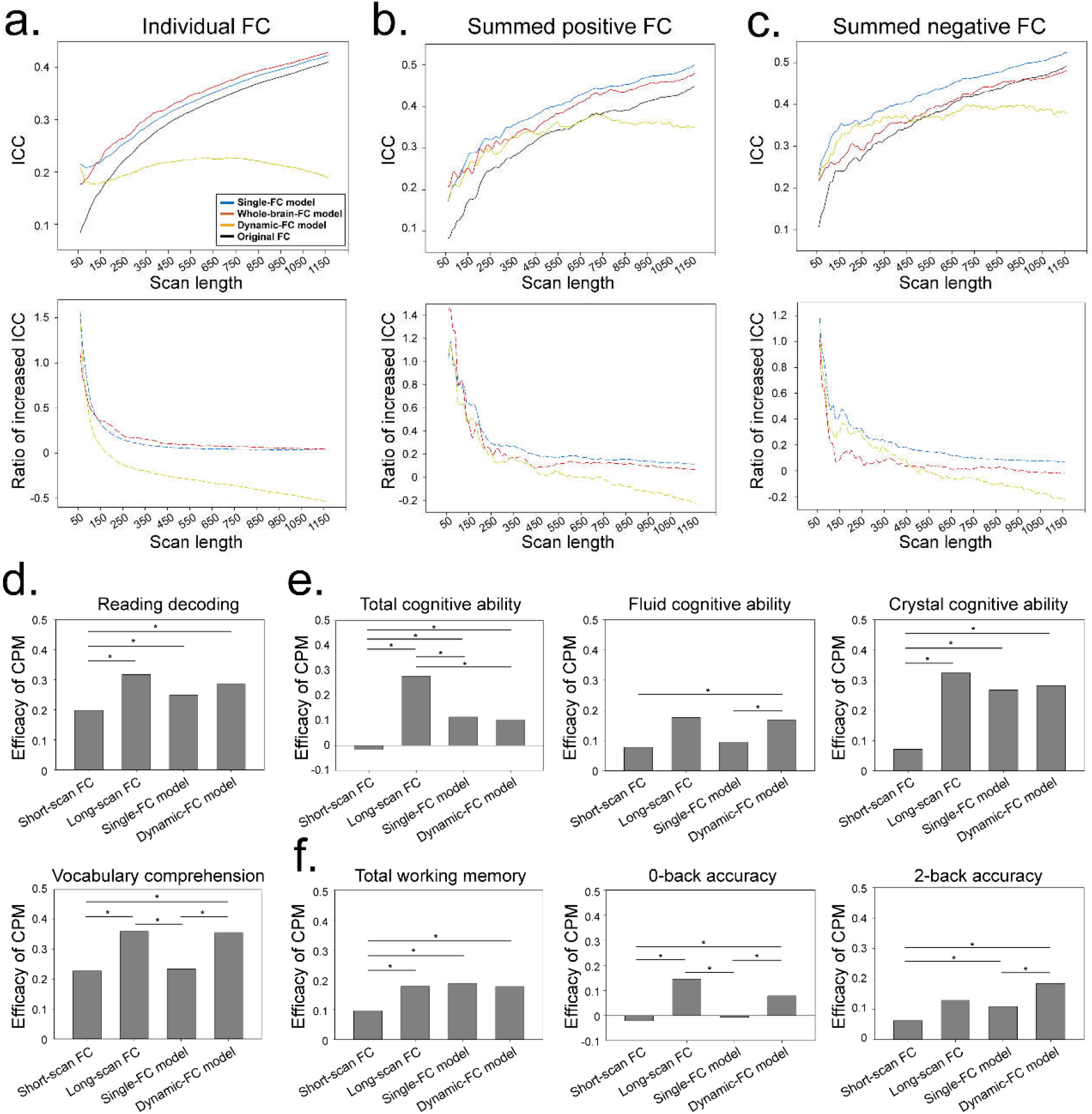
The application of predicted FC on intra-class correlation (ICC) and connectome-based predictive modeling (CPM). (a) Mean ICC of individual FC and the ratio of increase. (b) ICC of summed positive FC and the ratio of increase. (c) ICC of summed negative FC and the ratio of increase. (d)The efficacy of CPM to predict language ability. (e)The efficacy of CPM to predict cognitive ability. (f)The efficacy of CPM to predict working memory. Asterisk indicated significant difference (P<0.05 after Bonferroni correction).

The efficacy of CPMs based on long-scan FC and predicted FC (single-FC model and dynamic-FC model) were significantly higher than those based on the short-scan FC (**Fig 5d-5f and Table S2**). In most cases, the efficacy of CPMs based on long-scan FC was not significantly different from predicted FC. The performance of whole-brain-FC model was poor and not displayed because there were too many FCs than subjects.

## 4. Discussion

In this study, we developed three GLMs to predict long-scan FC based on short-scan FC. We developed three GLM models that can be used to predict long-scan FC from short-scan FC, and these models were robust across different data sources and can be applied to improve the test-retest reliability of FC and the performance of CPM.

### 4.1 The reliability of FC

In the last decade, many enlightening findings about the reliability of FC were reported: (1) static FC was more reliable than dynamic FC (26); (2) FC collected in eye-open condition was more reliable than eye-closed condition (6); (3) FC based on longer fMRI scan was more reliable (7); (4) FC based on concatenating fMRI data from multiple short runs was more reliable than a single long run (11); (5) FCs in high-order association regions, especially the frontal-parietal network and the default mode network, was more reliable (26, 27); (6) controversial were exist in whether FC in sensorimotor and visual networks was more reliable (26, 27); (7) reliability of dynamic FC could be improved in naturalistic viewing condition compared to that in resting state (28); (8) during movie watching, reliability in the primary visual cortex was decreased, but in higher-order regions was increased (27, 28). But to the best of our knowledge, our study is the first to increase the reliability of FC by predicting long-scan FC based on short-scan FC.

### 4.2 The potential application of models

Our models performed better when the scan length was shorter, suggesting their potential in enhancing the reliability of functional connectivity (FC), particularly for FC with extremely short scan lengths. Additionally, collecting extreme-long-scan data to train the model, such as the Midnight Scan Club (29), may be a viable method to increase the reliability of FC with normal scan length. However, in the leave-one-site-out validation, some datasets showed poor results, suggesting that inherent differences between datasets may still exist. Therefore, researchers may need to collect new data with the same parameters as before to apply to their previous data.

### 4.3 Linear model in FC-based researches

Our models, which were primarily based on linear models, outperformed several machine learning models, including support vector machines and LASSO models (**Table S3**). This could be attributed to the strong linear relationship between short-scan FC and long-scan FC, or the fact that the data size was still insufficient to train machine learning models. Additionally, studies had also shown that linear models were robust in predicting behavioral features (30), such as attention function (31), PTSD symptom prediction (32), autism (33), and so on. It is unlikely that this was due to the linear drift of the noise while scanning, as the process of removing the linear trend noise did not affect the results.

### 4.4 Limitations

Our research had some limitations. Firstly, we focused on predicting the resting state long-scan FC and did not investigate the dynamic features of long-scan FC, which requires further research. Secondly, the effect size was medium, and future studies could increase the effect size by using more advanced algorithms. Thirdly, the phase of dynamic FC may vary from different subjects in the dynamic-FC model, but this variation could not be captured by the GLM model.

## 5. Conclusion

We developed three GLM models that can be used to predict long-scan FC from short-scan FC, and these models were robust across different data sources and can be applied to improve the test-retest reliability of FC and the performance of CPM.

## Supporting information

Supplementary materials and Results

## Acknowledgements

Our gratitude goes for Dr. Wu-Xun Cui, Si-Jie Xiu from Department of Radiology of Tangdu Hospital, and for Dr. Kai Ai from department of Clinical Science of Philips Healthcare for their outstanding technique support.

