## Supplementary materials and Results for "Increasing the reliability of functional connectivity by predicting long-scan functional connectivity based on short-scan functional connectivity: model exploration, explanation, validation, and application"

### 1. Supplementary methods

#### 1.1 Subjects

Three datasets were used in this research, including the HCP dataset (N = 1003), the CoRR dataset (N = 904 from 18 sites), and our local dataset (N = 38).

##### 1.1.1 HCP dataset

The HCP began in 2010 with the goal of obtaining a dataset of unprecedented size and quality for mapping the normal human macro-scale connectome (1). The HCP young adults dataset was used (2017 release), which included 1003 subjects that aged between 22 and 35, and all subjects completed 4 runs of fMRI scan (14.4 mins per run).

##### 1.1.2 CoRR dataset

The goal of CoRR is to create an open scientific resource for the imaging community that helps to assess the test-retest reliability and reproducibility of functional and structural connectomics (2). CoRR collects previously collected test-retest imaging datasets from more than 36 laboratories around the world. To improve study quality, 904 subjects with similar scan parameters from 18 sites were selected from all 1629 subjects (described in *Imaging data*), all of whom completed at least 2 runs of fMRI scan (at least 5 mins per run).

##### 1.1.3 Local data

Our local data (3, 4) included 38 subjects that aged between 18-22, and all subjects completed 2 runs of fMRI scan (6.2 mins per run). To increase the consistency with CoRR dataset, the later 1.2 mins fMRI scan was deleted to obtain a 5-mins fMRI scan.

#### 1.2 Imaging data

##### 1.2.1 HCP dataset

The time series of all brain nodes of all subjects that extracted from the minimally pre-processed fMRI data (21) were directly downloaded and used (https://db.humanconnectome.org/data/projects/HCP_1200). Briefly speaking, all subjects have four runs of 14.4-min multi-band accelerated resting-state fMRI scans with a 0.72-second time of repetitions (TR), thus the total scan length was 4800 TRs (4 run×14.4 min/run×60 s/min×1/0.72 TR/s). All subjects were scanned with 3T Siemens scanners, with an isotropic spatial resolution of 2 mm. Pre-processing steps included corrections for spatial distortions and head motion, registration to the T1 weighted structural image, resampling to 2 mm MNI space, and the FIX artefact removal procedure was used to denoise. Group independent component analysis was used to parcellate the whole brain into different (15, 25, 50, 100, 200, 300) independent components (ICs). For a given parcellation, dual-regression was used to estimate the nodal timeseries. In this procedure, the full set of ICA maps was used as spatial regressors against the full data, estimating one representative timeseries for each ICA map for each subject. Finally, 1003 (subjects)×4800 (scan length)×N (ICs) timeseries were obtained.

#### 1.2.2 CoRR dataset

Different TRs can lead to a discrepancy between scan length and the scan time (for example, when TR is 3 seconds, the scan length obtained by a 6-minutes scan is 120 TR, but when TR is 2 seconds, the scan length is 180 TR). In order to improve the quality of research, only datasets with a TR of 2 seconds were selected, including 904 subjects from 18 sites. Subjects in CoRR dataset were all aged between 7 and 84 and completed at least 2 runs of fMRI scan (at least 5 mins per run), thus the total scan length was at least 300 TRs (2 run×5 min/run×60 s/min×1/2 TR/s). After that, traditional pre-processing and denoising procedures were applied by using Statistical Parameter Mapping (SPM12, Wellcome, Imaging Neurology Group, London, UK; http://www.fil.ion.ucl.ac.uk/spm) and DPABI toolkit (5), including slice timing, correction of head motion, registration to the T1 weighted structural image, resampling to 2 mm MNI space, smooth with a 6 mm full width at half maximum (FWHM) Gaussian kernel, nuisance signals regression (including the 24-parameter head motion profile, white matter, and cerebrospinal fluid signals), and band-pass filtering with a 0.01~0.1 Hz band. The coordinates of peak values of ICs in HCP dataset (15-IC parcellation) were defined as the region of interest (ROI), and time series were extracted by averaging the value around ROI (radius = 6 mm). Finally, 904×300×15 timeseries were obtained.

##### 1.2.3 Local data

The procedures were the same as CoRR dataset. Finally, 38×300×15 timeseries were obtained.

#### 1.3 Study design

This research included four key experiments: model exploration, explanation, validation, and application. In model exploration step, three GLM models were established to predict the long-scan FC based on the short-scan FC. In model explanation step, we explained these models by visualization of weights in above models and analyzation of distribution features of predicted FC matrices. In model validation step, our results were validated in different datasets, including CoRR project and our local dataset. In model application step, these models were applicated to improve the test-retest reliability of FC and the performance of connectome-based predictive models (CPM).

#### 1.4 Model exploration: developing three models to predict long-scan FC based on short-scan FC

##### 1.4.1 Models

The first model directly used the short-scan FC to predict its long-scan form, which was called single-FC model:

$$Corr(T_i^{[1,n]}, T_j^{[1,n]}) = \beta_1 Corr(T_i^{[1,t]}, T_j^{[1,t]}) + \beta_2 A + \beta_3 S + e;$$

Where *T* was the timeseries, *i* and *j* were corresponding ICs of the FC, *n* was the scan length of the long-scan timeseries, *t* was the scan length of the short-scan timeseries, *A* was age, *S* was sex, and *e* was error.

The second model used whole-brain FCs to predict each long-scan FC, which was called whole-brain-FC model:

$$Corr(T_i^{[1,n]}, T_j^{[1,n]}) = \sum_{a=1}^{IC}\sum_{b=1}^{IC} \beta_{(i,j)}\, Corr(T_a^{[1,t]}, T_b^{[1,t]}) + \beta_{IC\times\frac{IC-1}{2}+1} A + \beta_{IC\times\frac{IC-1}{2}+2} S + e;$$

Where *T* was the timeseries, *i* and *j* were corresponding ICs of the FC, *n* was the scan length of the long-scan timeseries, *t* was the scan length of the short-scan timeseries, *a* and *b* were ICs, *IC* was the number of ICs, *A* was age, *S* was sex, and *e* was error.

The third model used a sliding-window method to extract the dynamic FC of single FC to predict its long-scan form, which was called dynamic-FC model:

$$Corr(T_i^n, T_j^n) = \sum_{k=1}^{(t-w+1)/s} \beta_k Corr(T_i^{[((k-1)\times s+1),((k-1)\times s+w)]}, T_j^{[((k-1)\times s+1),((k-1)\times s+w)]})$$

$$+ \beta_{(\frac{t-w+1}{s})+1} A + \beta_{(\frac{t-w+1}{s})+2} S + e;$$

Where *T* was the timeseries, *i* and *j* were corresponding ICs of the FC, *n* was the scan length of the long-scan timeseries, *t* was the scan length of the short-scan timeseries, *w* was the window width of dynamic FCs, *s* was the step length of dynamic FCs, *k* was the number of dynamic FCs, *A* was age, *S* was sex, and *e* was error. In this model, common window width (40 TRs) and step length (3 TRs) were used, while other conditions were also tested.

#### 1.4.2 Variables

Three variables may influence the result, including scan length, subject numbers, and methods of parcellation. Consequently, all conditions of each variable were tested while the condition of other two variables were fixed at the same time (**Fig 1c**).

**With scan length as variable:** The long-scan fMRI time series (4800 TRs) was correlated with each other to generate the long-scan FC. In the calculation of short-scan FC, the scan length was successively extended from 50 TRs to 4750 TRs, with an interval of 50 TRs. In this procedure, 603 subjects (~60%) were randomly selected as the training set and the rest 400 subjects (~40%) as the test set. Besides, the method that parcellated the brain into 15 ICs were used.

**With subject numbers as variable:** The number of subjects in the training set was successively decreased from 953 to 53, with an interval of 50 subjects, and the remaining subjects were the test set. In order to prevent bias caused by the selection of subjects, the selection procedure was randomly repeated 500 times, and the results were shown as the mean ± standard deviation of 500 repetitions. Since most current studies have scan time of around 6 minutes, short scan was defined as fMRI time series within the first half of the first run (600 TRs, about 7.2 minutes). Besides, the method that parcellated the brain into 15 ICs were used.

**With independent component numbers as variable:** The brain was parcellated into different Independent components (from 15 ICs to 300 ICs). In this procedure, short scan was defined as fMRI time series within the first half of the first run, and 603 subjects were randomly selected as the training set and the rest 400 subjects as the test set.

##### 1.4.3 Outcome assessments

The results were evaluated by comparing the absolute distance (**Fig 1e**) between long-scan FC and short-scan FC ($D_{|L-S|}$) with the absolute distance between long-scan FC and predicted FC ($D_{|L-P|}$). The model is considered effective if the long-scan FC is closer to predicted FC than to the short-scan FC ($D_{|L-P|} < D_{|L-S|}$).

#### 1.5 Model explanation: visualization of weights in above models and analyzation of distribution features of predicted FC matrices

The beta values in above three models were visualized to explain the contribution of each component in each model. Besides, the distribution features of FC across individual brain, including mean value, standard deviation, kurtosis, and skewness, were compared between three models.

#### 1.6 Model validation: validating models in CoRR and our local datasets

Two runs of fMRI time series (150 TRs per run) were concatenated together to form a long-scan fMRI time series (300 TRs). Then, the long-scan fMRI time series was correlated with each other to form the long-scan FC matrix. In the calculation of short-scan FC, the scan length was successively extended from 40 TRs to 290 TRs, with an interval of 10 TRs. Besides, the method that parcellated the brain into 15 ICs were used.

In the first step, all subjects were pooled together and randomly divided into the training set (565 subjects, 60%) and the test set (377 subjects, 40%). In the second step, a leave-one-site-out cross validation method was used, with each set was selected out as the test set, and the rest as training set in every time of model training.

#### 1.7 Model application: applying predicted FC matrix on the test-retest reliability and connectome-based predictive model

In the first step, the test-retest reliability was measured among the original FC values based on the four runs of fMRI scan as well as among the predicted FC values based on aforementioned three models. Then, the ratio of increase of test-retest reliability was calculated.

In the second step, the FC matrices (to obtain best performance, the method to parcellate the brain into 300 ICs was used) of 400 subjects in test set were used to predict language ability (including oral reading recognition and picture vocabulary), cognitive ability (including fluid, crystal, and total cognitive function composite), and working memory (N-back task).

#### 1.8 Statistical analysis

##### 1.8.1 Model exploration

**The shortened absolute FC distance of a single FC (SADS):** Effect size (Cohen's d) was calculated to qualify the shortened absolute FC distance from $D_{|L-S|}$ to $D_{|L-P|}$ (**Fig 1f**).

**The shortened absolute FC distance of the sum of all positive FCs (SADP):** All positive values of the short-scan, long-scan, and predicted FC matrices were respectively summed to generate a single value (**Fig 1f**). Then, $D_{|L-S|}$ was calculated by the absolute difference between the positive value of short-scan FC and long-scan FC, and $D_{|L-P|}$ was calculated by the absolute difference between the positive value of predicted FC and long-scan FCs.

**The shortened absolute FC distance of the sum of all negative FCs (SADN):** Similar to above, but all negative values in FC matrices were summed together.

##### 1.8.2 Model explanation

The mean value, standard deviation, kurtosis, and skewness of the FC matrix were calculated for each subject. Analysis of variance (ANOVA) or non-parametric Mann-Whitney U test was used (depending on whether it was object to the Gaussian distribution) to compare these four features between short-scan, long-scan, and predicted FC matrices.

##### 1.8.3 Model validation

All parameters were the same as *Model exploration* step.

##### 1.8.4 Model application

**Test-retest reliability:** The intraclass correlation coefficient (ICC) was calculated to represent the test-retest reliability among FC values and summed FC values based on four sessions of fMRI scan.

**Efficacy of CPM:** Leave-one-out cross-validation was used for model training (32). First, one subject was leaved out as test set, and the other subjects were used as training set. Second, for all subjects in the training set, each edge in the connectivity matrix was associated with the behavior performance (either Pearson correlation or Spearman correlation, depending on whether the data is normally distributed). Third, the strongest FC edges (above the significance threshold of 0.001) were selected for further correlation analysis. Fourth, the selected edges were separate into positive and negative groups, and both groups of edges were summed together to form 2 values for each subject. fifth, a linear model was built with the summed values and behavior variables as independent and dependent variables, separately. Finally, the leaved out subject was used to test the efficacy of the model. In this step, the connectivity matrix mask obtained in the training procedure was applied on the testing subject. When all procedures were completed, there was a predicted value for each subject, and correlation analysis between predicted values and true values were used to reflect the efficacy of CPMs.

### 2. Supplementary results

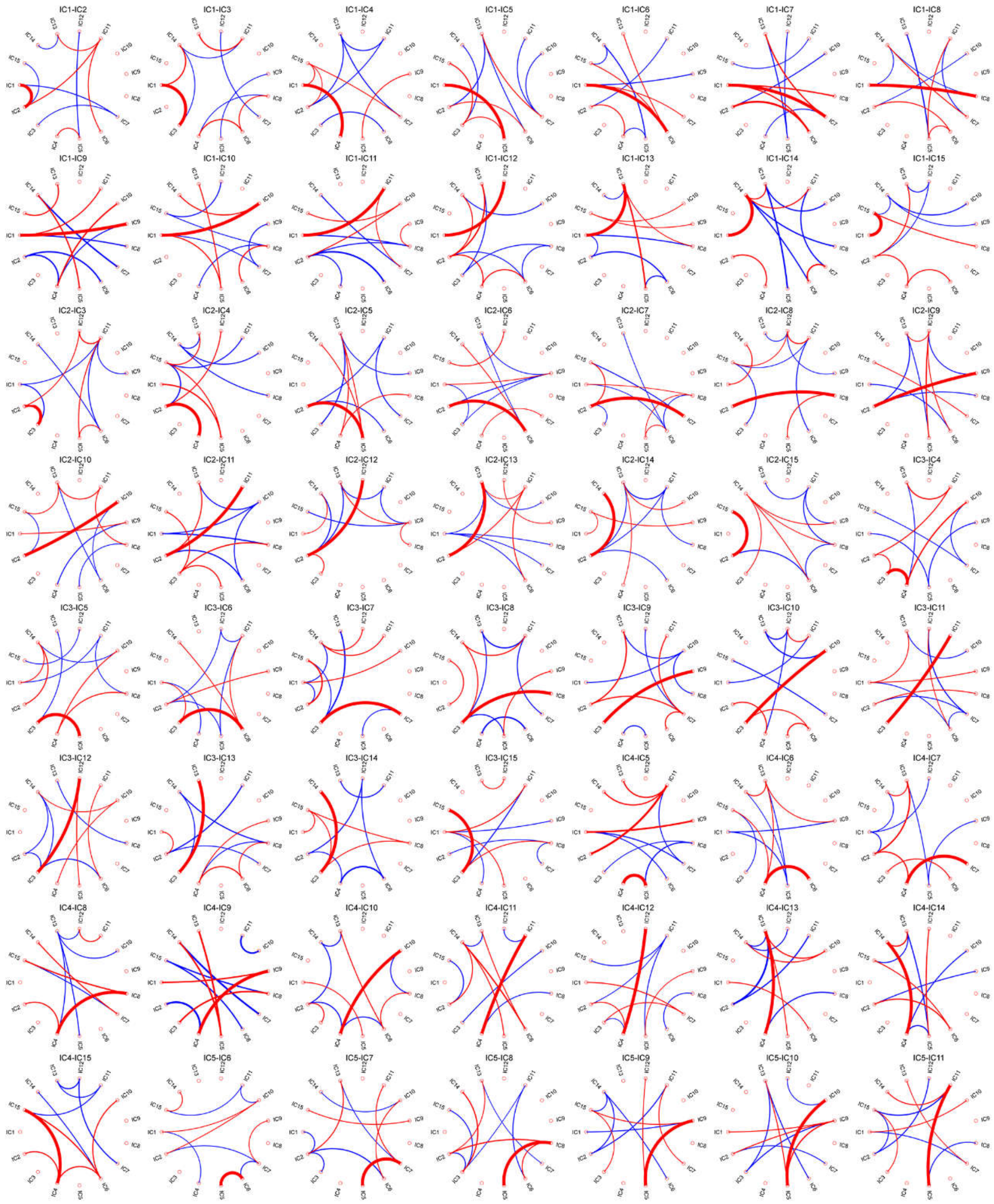

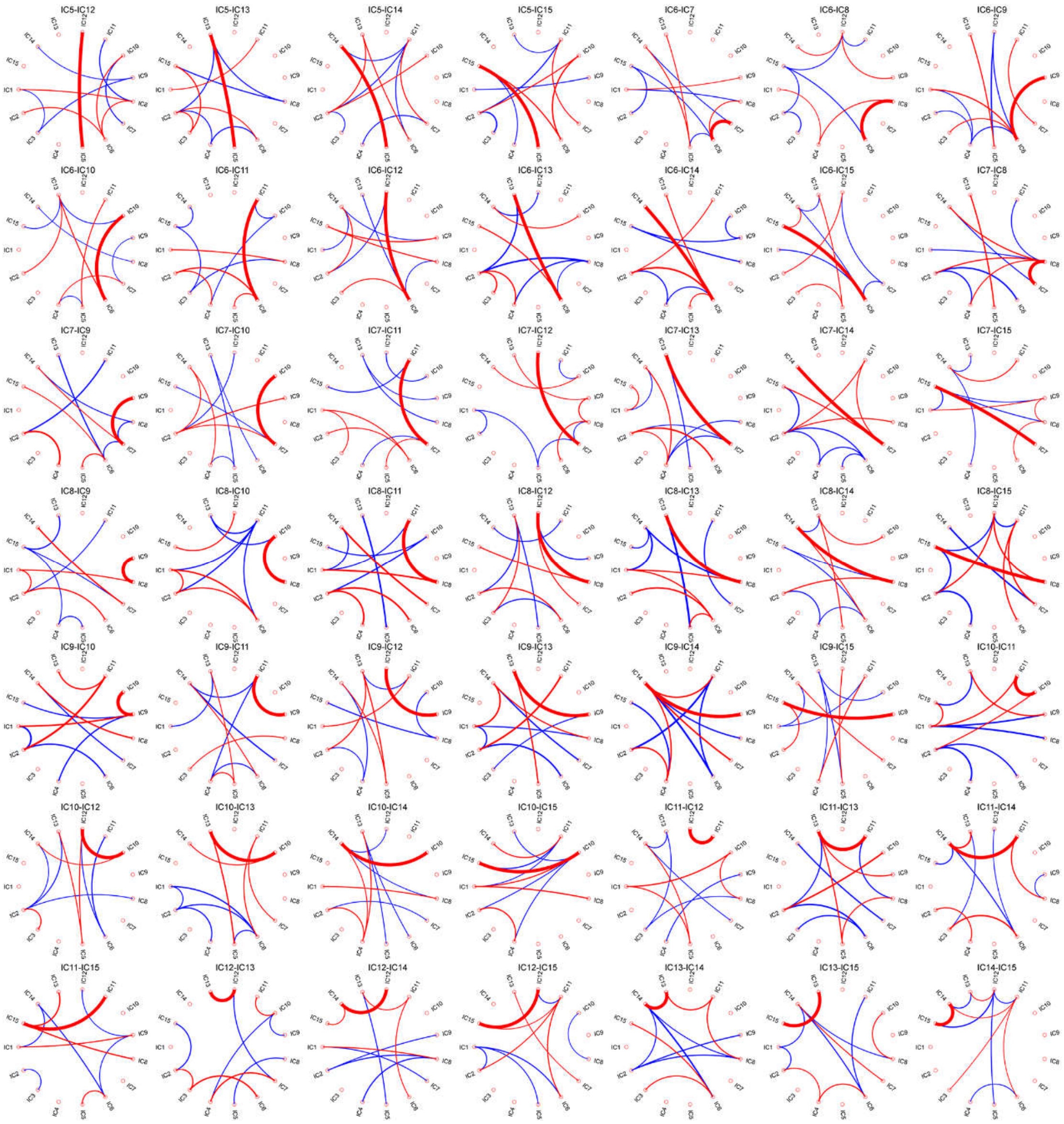


**Fig S1. The strongest weights (top 10) in the whole-brain-FC model to predict the long-scan FC between each two ICs.**

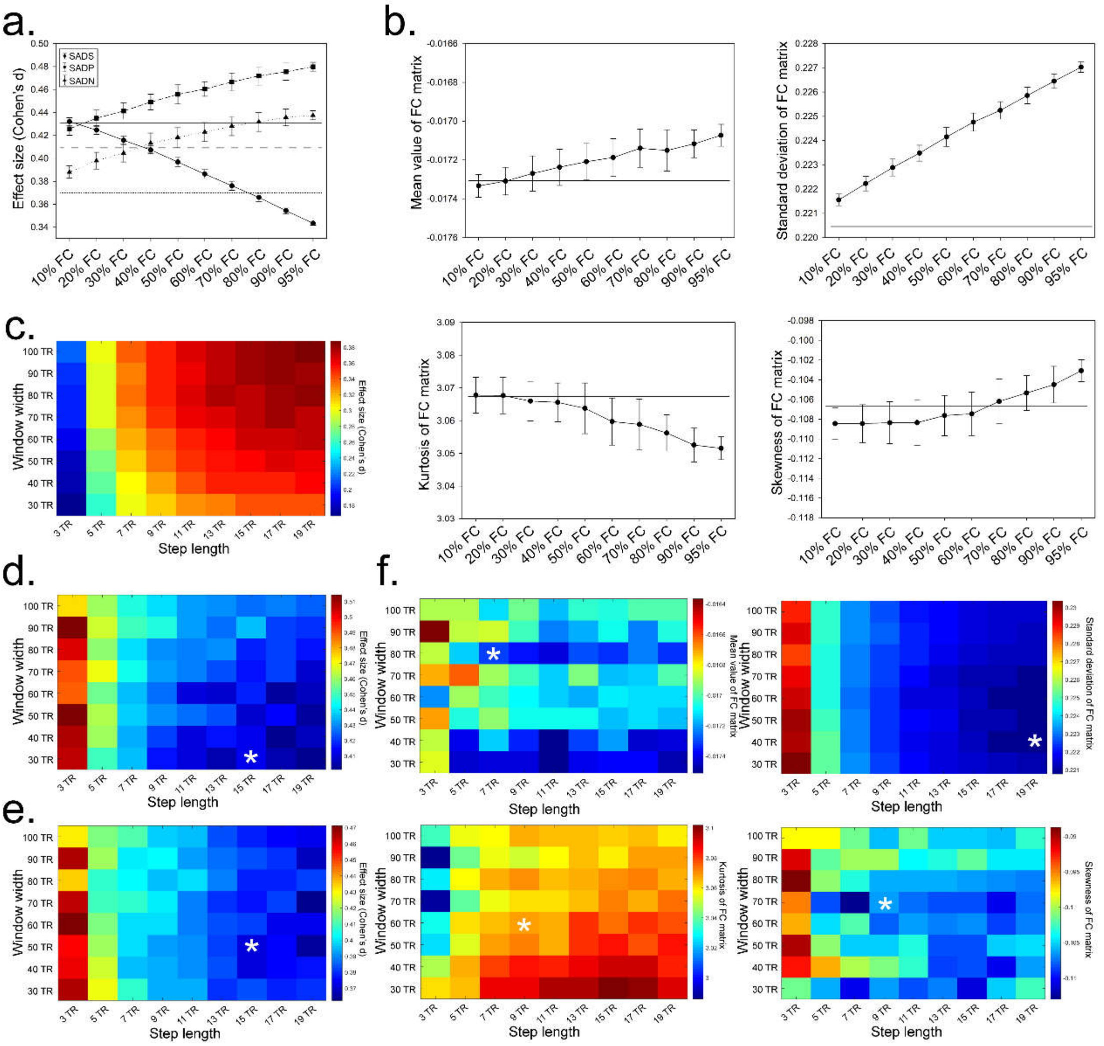


In this study, we aimed to investigate the tradeoff between the number of FCs included in the model and the results obtained. We compared a single-FC model, which included only one FC, with a whole-brain-FC model, which included all FCs. To test the interim condition, we successively selected FCs from 10% to 90% and analyzed the changes in SADS, SADP, SADN, and distribution features. Each selection of FC was randomly conducted and repeated 100 times, and the results were presented as the mean ± standard deviation of 100 repetitions. Additionally, we used a traditional window width (40 TRs) and a traditional step length (3 TRs) in the dynamic-FC model. We also tested the results using other window widths and step lengths.

**Fig S2. The tradeoff between FC numbers that included in model and results.** (a) When the FC numbers that included in whole-brain-FC model gradually increased, the SADS gradually decreased and the SADP and SADN gradually increased. (b) When the FC numbers that included in model gradually increased, the distribution of predicted FC matrix became more discrete, manifested as increased standard deviation and decreased kurtosis. (c) The changes of SADS along with the changes of window width and step length in the calculation of dynamic FC. (d) The changes of SADP along with the changes of window width and step length in the calculation of dynamic FC. (e) The changes of SADN along with the changes of window width and step length in the calculation of dynamic FC. (f) The changes of distribution features of predicted FC matrix along with the changes of window width and step length in the calculation of dynamic FC. Asterisk mean the value equal to that derived from sing-FC model.

**Table S1. Demographic information and parameters of fMRI scan.**

| Data sources | Subject number (Included/Total) | Age (mean±std) | Sex (female/male) | Manufacturer | Field Strength (T) | TR (s) | Number of slices | Included runs | Max scan length per run |
|---|---|---|---|---|---|---|---|---|---|
| HCP project | 1003/1003 | 28.7±3.7 | 534/469 | Siemens | 3 | 0.72 | 72 | 4 | 1200 |
| CoRR project | | | | | | | | | |
| BNU1 | 51/57 | 23.10±2.39 | 26/25 | Siemens | 3 | 2 | 33 | 2 | 200 |
| BNU3 | 48/48 | 22.54±2.15 | 24/24 | Siemens | 3 | 2 | 34 | 2 | 150 |
| HUN1 | 30/30 | 24.37±2.41 | 15/15 | GE | 3 | 2 | 43 | 2 | 300 |
| IPCAS1 | 30/30 | 20.90±1.77 | 9/21 | Siemens | 3 | 2 | 32 | 2 | 205 |
| IPCAS3 | 28/36 | 21.18±1.89 | 9/19 | Siemens | 3 | 2 | 64 | 2 | 180 |
| IPCAS4 | 20/20 | 23.15±1.60 | 10/10 | GE | 3 | 2 | 37 | 2 | 180 |
| IPCAS5 | 21/22 | 18.29±0.46 | 21/0 | Siemens | 3 | 2 | 33 | 2 | 170 |
| IPCAS8 | 13/13 | 57.62±3.78 | 6/7 | Siemens | 3 | 2 | 33 | 2 | 240 |
| JHNU1 | 30/30 | 23.27±3.69 | 21/9 | Siemens | 3 | 2 | 30 | 2 | 250 |
| MRN1 | 38/56 | 21.87±9.28 | 20/18 | Siemens | 3 | 2 | 33 | 2 | 150 |
| NYU1 | 25/25 | 29.44±8.64 | 10/15 | Siemens | 3 | 2 | 33 | 2 | 180 |
| NYU2 | 186/187 | 20.23±11.49 | 115/71 | Siemens | 3 | 2 | 33 | 2 | 180 |
| SWU1 | 20/20 | 21.55±1.76 | 6/14 | Unknow | 3 | 2 | 33 | 2 | 240 |
| SWU2 | 27/27 | 20.96±1.65 | 9/18 | Unknow | 3 | 2 | 32 | 2 | 242 |
| SWU3 | 23/24 | 20.43±1.65 | 8/15 | Unknow | 3 | 2 | 32 | 2 | 300 |
| SWU4 | 225/235 | 20.04±1.27 | 118/107 | Unknow | 3 | 2 | 32 | 2 | 242 |
| UM | 69/80 | 65.55±6.26 | 22/47 | Unknow | 3 | 2 | 32 | 2 | 150 |
| UTAH1 | 20/26 | 18.85±7.93 | 20/0 | Siemens | 3 | 2 | 40 | 2 | 240 |
| Local data | 38/42 | 20.76±0.97 | 0/38 | GE | 3 | 2 | 36 | 2 | 150 |

Images that were distorted after normalization were excluded.

**Table S2. The comparison of efficacy of CPM.**

| Efficacy of CPM | Short-scan FC | Long-scan FC | Single-FC model | Dynamic FC model | Comparison |
|---|---|---|---|---|---|
| Reading decoding | 0.1982 | 0.3177 | 0.2490 | 0.2854 | |
| P value (Corrected) | 0.0066 | | | | Long-scan FC |
| P value (Corrected) | <0.001 | 0.0773 | | | Single-FC model |
| P value (Corrected) | 0.0017 | 0.2610 | 0.0970 | | Dynamic FC model |
| Vocabulary comprehension | 0.2278 | 0.3603 | 0.2346 | 0.3567 | |
| P value (Corrected) | 0.0012 | | | | Long-scan FC |
| P value (Corrected) | 0.3416 | 0.0022 | | | Single-FC model |
| P value (Corrected) | <0.001 | 0.4666 | <0.001 | | Dynamic FC model |
| Total cognitive ability | 0.0776 | 0.1763 | 0.0951 | 0.1680 | |
| P value (Corrected) | 0.0092 | | | | Long-scan FC |
| P value (Corrected) | 0.0401 | 0.0263 | | | Single-FC model |
| P value (Corrected) | <0.001 | 0.4218 | <0.001 | | Dynamic FC model |
| Fluid cognitive ability | -0.0167 | 0.2786 | 0.1143 | 0.1008 | |
| P value (Corrected) | <0.001 | | | | Long-scan FC |
| P value (Corrected) | <0.001 | <0.001 | | | Single-FC model |
| P value (Corrected) | <0.001 | <0.001 | 0.3169 | | Dynamic FC model |
| Crystal cognitive ability | 0.0716 | 0.3238 | 0.2671 | 0.2823 | |
| P value (Corrected) | <0.001 | | | | Long-scan FC |
| P value (Corrected) | <0.001 | 0.1043 | | | Single-FC model |
| P value (Corrected) | <0.001 | 0.2041 | 0.3175 | | Dynamic FC model |
| Total working memory | 0.0970 | 0.1778 | 0.1881 | 0.1772 | |
| P value (Corrected) | 0.0207 | | | | Long-scan FC |
| P value (Corrected) | <0.001 | 0.3955 | | | Single-FC model |
| P value (Corrected) | <0.001 | 0.4950 | 0.2240 | | Dynamic FC model |
| 0-back accuracy | 0.062646 | 0.1296 | 0.1091 | 0.1847 | |
| P value (Corrected) | 0.1025 | | | | Long-scan FC |
| P value (Corrected) | <0.001 | 0.3494 | | | Single-FC model |
| P value (Corrected) | <0.001 | 0.1497 | <0.001 | | Dynamic FC model |
| 2-back accuracy | -0.0217 | 0.1445 | -0.0068 | 0.0780 | |
| P value (Corrected) | <0.001 | | | | Long-scan FC |
| P value (Corrected) | 0.2559 | <0.001 | | | Single-FC model |
| P value (Corrected) | <0.001 | 0.0629 | <0.001 | | Dynamic FC model |

**Table S3. The efficacy of other machine learning models.**

| | SADS | SADP | SADN |
|---|---|---|---|
| Support vector machine | 0.418 | 0.442 | 0.400 |
| LASSO | -0.108 | -1.456 | -1.991 |
| Single-FC model | 0.433 | 0.413 | 0.372 |
| Whole-brain-FC model | 0.413 | 0.481 | 0.509 |
| Dynamic-FC model | 0.372 | 0.439 | 0.458 |

Scan length = 600 TRs; Subjects in training set = 603, subjects in test set = 400; IC numbers = 15.
